# Genome of the bi-annual Rio Pearlfish (*Nematolebias whitei*), a killifish model species for Eco-Evo-Devo

**DOI:** 10.1101/2021.11.24.469934

**Authors:** Andrew W. Thompson, Harrison Wojtas, Myles Davoll, Ingo Braasch

**Affiliations:** Department of Integrative Biology, Michigan State University, East Lansing, MI, USA; Ecology, Evolution & Behavior (EEB) Program, Michigan State University, East Lansing, MI, USA; Department of Biology, University of Virginia, Charlottesville, VA, USA

**Keywords:** diapause, aging, hatching, extreme environments, Eco-Evo-Devo, teleost

## Abstract

The Rio Pearlfish *Nematolebias whitei* is a bi-annual killifish species inhabiting seasonal pools of the Rio de Janeiro region that dry twice per year. Embryos enter dormant diapause stages in the soil, waiting for the inundation of the habitat which triggers hatching and commencement of a new life cycle. This species represents a convergent, independent origin of annualism from other emerging killifish model species. While some transcriptomic datasets are available for Rio Pearlfish, thus far a sequenced genome has been unavailable. Here we present a high quality, 1.2Gb chromosome-level genome assembly, genome annotations and a comparative genomic investigation of the Rio Pearlfish as representative of a vertebrate clade that evolved environmentally-cued hatching. We show conservation of 3-D genome structure across teleost fish evolution, developmental stages, tissues and cell types. Our analysis of mobile DNA shows that Rio Pearlfish, like other annual killifishes, possesses an expanded transposable element profile with implications for rapid aging and adaptation to harsh conditions. We use the Rio Pearlfish genome to identify its hatching enzyme gene repertoire and the location of the hatching gland, a key first step in understanding the developmental genetic control of hatching. The Rio Pearlfish genome expands the comparative genomic toolkit available to study convergent origins of seasonal life histories, diapause, and rapid aging phenotypes. We present the first set of genomic resources for this emerging model organism, critical for future functional genetic and multi-omic explorations of “Eco-Evo-Devo” phenotypes in resilience and adaptation to extreme environments.

**Significance:**

Seasonal or annual killifishes are emerging models for aging, life history adaptions to extreme environments, and ecological evolutionary developmental biology (Eco-Evo-Devo). Most studies have thus far focused on the African turquoise killifish *Nothobranchius furzeri* and the South American *Austrofundulus limneaus.* We sequenced and analyzed the genome of the Rio Pearlfish *Nematolebias whitei* from the Rio de Janeiro region, a seasonal species representing a convergent origin of seasonality from other sequenced killifish species, strengthening the comparative potential of Aplocheiloid killifishes as a model clade for the comparative and functional genomics of animal resilience to environmental change.

## Introduction

Aplocheiloid killifishes inhabit tropical freshwater habitats around the world. Some species in Africa and in the neotropics have evolved to live in ephemeral water bodies that are subject to seasonal cycles of desiccation and inundation (Simpson 1979; Myers 1952). Desiccation results in the death of the adult populations of these annual or seasonal killifishes. Annual killifishes show rapid aging due to relaxed selection on lifespan (Cui et al. 2019) and are an important emerging senescence model system with several sequenced genomes and amenability to functional genetic manipulation (Valenzano et al. 2011; Reichwald et al. 2015; Harel et al. 2015; Valenzano et al. 2015; Wagner et al. 2018). Annual killifish embryos survive the dry season encased in specialized eggs (Thompson et al. 2017) buried in the soil by undergoing three distinct diapause stages (DI, DII, DIII; Wourms 1972a, 1972b, 1972c). DI occurs as a migratory dispersion of blastomeres that later coalesce to commence development. DII occurs during somitogenesis when organs are rudimentary, and DIII occurs after organogenesis when the embryo is fully formed, poised to hatch and then immediately feed as free-swimming larva upon inundation of the habitat. The seasonal life history of annual killifishes is a remarkable example of convergent evolution with up to seven losses or gains across aplocheiloid killifish evolution (Thompson et al. 2021).

The Rio Pearlfish *Nematolebias whitei* (Cyprinodontiformes: Rivulidae) is a seasonal killifish endemic to the coastal plains of the Rio de Janeiro region in Brazil, inhabiting small pools that dry twice annually, usually from July to August and February to March (Fig. 1A, Myers 1942; Costa 2002). Rio Pearlfish represents a separate origin of seasonality from the other seasonal killifish model species *Nothobranchius furzeri* and *Austrofundulus limnaeus* (Thompson et al. 2021; Furness et al. 2015; Reichwald et al. 2015; Valenzano et al. 2015). In *N. whitei*, DI and DII are facultative and DIII can be a “prolonged”, “deep” stasis compared to other diapausing killifishes, occurring just before environmentally-cued hatching upon submersion in water (Wourms 1972c; Thompson & Ortí 2016). Rio Pearlfish was suggested as a top candidate species for annual killifish research model systems in the seminal work of developmental biologist John P. Wourms in 1967. They are small-bodied, prolific, and hardy, and readily spawn in silica sand, making them easily-maintained laboratory animals (Wourms 1967) and amenable to genetic manipulation similar to other killifish species (Harel et al. 2015; Aluru et al. 2015). Rio Pearlfish has also been an emergent system to study aging (Ruijter 1987), environmental influences on development (Ruijter et al. 1984), the role of prolactin in hatching control (Schoots et al. 1983; Ruijter & Creuwels 1988), resilience to perturbations in development with the ability to develop normally from diblastomeric eggs (Carter & Wourms 1993), and more recently, the transcriptional control of diapause and hatching (Thompson & Ortí 2016). These studies provide important insights into suspended animation, environmentally cued hatching, resilience to extreme environmental conditions, and developmental abnormalities with important implications for biomedical studies.

**Figure 1.**
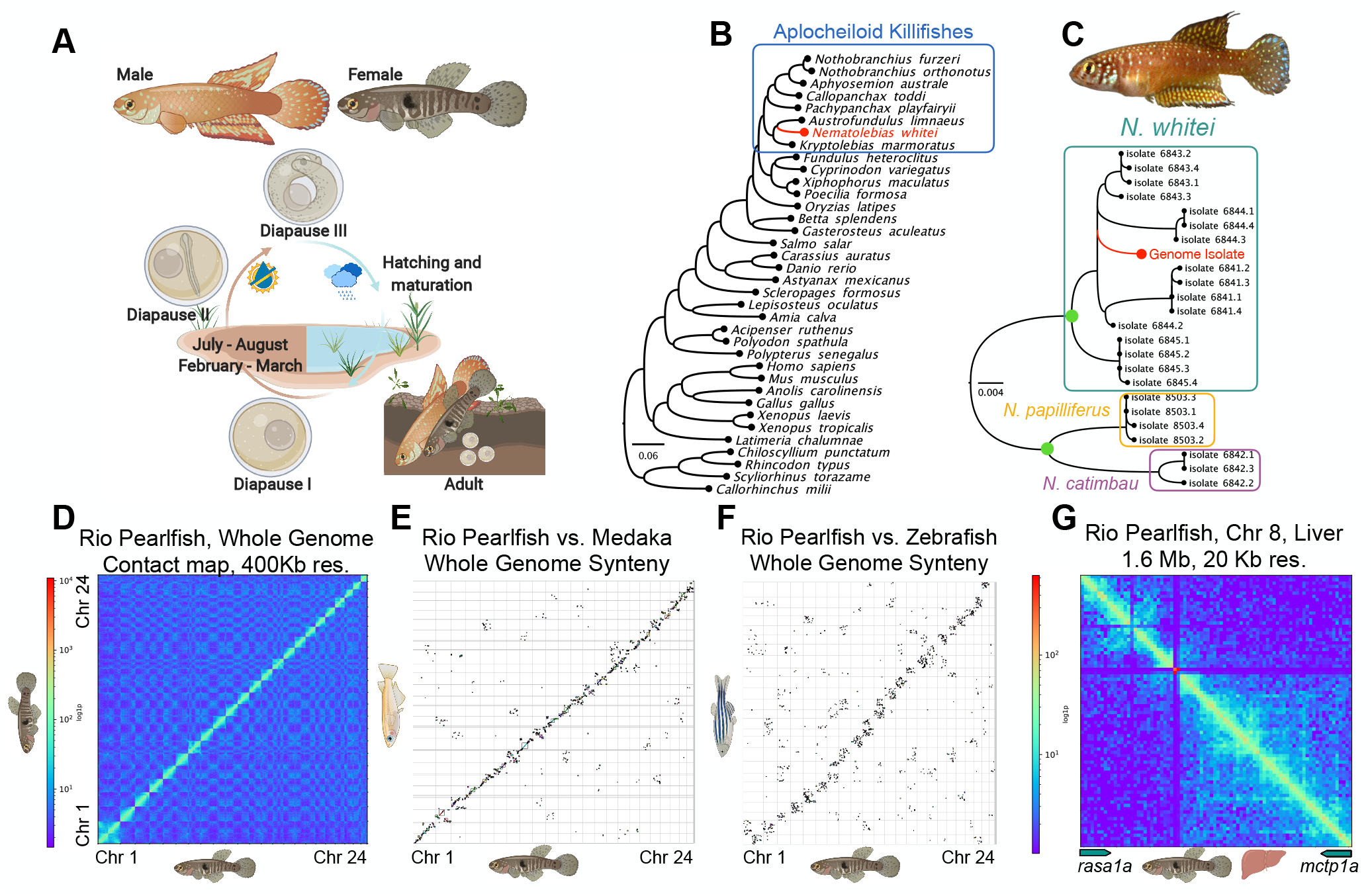
Rio Pearlfish evolution, ecology, development, and 3-D genome structure. A.) Bi-annual life cycle of the Rio Pearlfish with three developmental diapause stages upon burying eggs in soil. B.) Relative position of Rio Pearlfish in the vertebrate tree of life inferred by Orthofinder2 and based annotated proteins. C.) DNA barcode (*cox1* and *cytb*) phylogeny inferred with RAxML of the genus *Nematolebias* confirming the identity of the genome specimen as *N. whitei.* Sequences from Costa et al. (2014) were used for comparison to the genome sequence. Green nodes show 100% bootstrap support for the reciprocal monophyly of *N. whitei* with other genera and confirms the identity of the genome specimen with high confidence. D.) Hi-C contact map of the Rio Pearlfish genome showing linkage of the 24 chromosomes into chromosomal pseudomolecules. E-F.) SynMap genome-wide synteny plots of Rio Pearlfish vs. medaka (E) and vs. zebrafish (F) generated showing high conservation across over 250 million years of teleost evolution. G.) Hi-C contact maps of the syntenic region between *rasa1a* and *mctp1a* in pearlfish liver tissue. These contact maps highlight the conserved 3-D structure that include topologically associated domains (TADs) conserved across teleost evolution as well as cell types and developmental stages (Nakamura et al. 2021). Species graphics generated with BioRender.

A high-quality reference genome sequence and annotation as well as a characterization of gene orthology and gene synteny are critical for understanding how evolutionary genomic changes generate biodiversity. New tools for investigation of 3-D genome structure and the epigenome inform studies on gene synteny, chromosomal rearrangements, and the control of progression from genotype to phenotype. For example, three-dimensional chromatin structure impacts gene regulation and can manifest as topologically associated domains (TADs) with high frequencies of internal physical self-contacts that could represent higher order gene regulatory regions conserved across evolutionary timescales (Krefting et al. 2018). However, 3-D genome structure remains uncharacterized in annual killifishes.

Likewise, mobile genetic elements can have far-reaching effects on genotype and phenotype. Transposable elements (TEs) are hypothesized to generate novel genetic substrate for adaptations (Casacuberta & González 2013; Esnault et al. 2019; Feiner 2016). Some annual fishes have substantially expanded TE content compared to their non-annual relatives (Cui et al. 2019), and the link between TEs, aging, and human disease phenotypes (Bravo et al. 2020) coupled with the rapid senescence of annual killifishes highlights the importance of examining the “mobilome” in emerging research organisms like the Rio Pearlfish and other killifishes.

Hatching is a fundamental process during animal develepment, and aquatic vertebrates hatch via release of choriolytic enzymes secreted from hatching gland cells (Hong & Saint-Jeannet 2014; Yamagami 1988) that break down the egg envelope or chorion. The common ancestor of clupeocephalan teleost fishes underwent a hatching enzyme gene duplication followed by divergence and neo/subfunctionalization into the high choriolytic enzyme (*hce*) and low chorioltyic enzyme genes (*lce*) (Yasumasu et al. 1992; Kawaguchi et al. 2010, 2006; Sano et al. 2014). However, the location and development of hatching gland cells (HGCs) are dynamic among teleost fishes (Inohaya, et al. 1995; Inohaya et al. 1997) and migrate and localize in different anatomical positions in different species at the time of hatching (Shimizu et al. 2014; Nagasawa et al. 2016; Korwin-Kossakowski 2012). Pinpointing HGC location in seasonal killifishes is necessary for understanding the genetic and environmental regulation of their highly orchestrated hatching process under extreme environmental conditions.

Through genomic studies on Rio Pearlfish we hope to expand the fish “model army” (Braasch et al. 2015) and the killifish “model clade” (Sanger & Rajakumar 2019) as a comparative approach is critical for understanding the evolution of complex developmental and aging phenotypes. Here, we construct a chromosome-level genome assembly for the Rio Pearlfish, utilizing Hi-C contact maps, genome annotations, and gene expression analyses to characterize genomic evolution and hatching biology in these extremophilic fishes. These genomic resources make Rio Pearlfish a tractable model for genetic exploration study of their unique “Eco-Evo-Devo” phenotypes.

## Materials and Methods

### Genome Sequencing and Assembly

A total of 1.25 ng of template gDNA extracted from the liver of a single adult female *N. whitei* was loaded on a Chromium Genome Chip. Whole genome sequencing libraries were prepared using 10X Genomics Chromium Genome Library & Gel Bead Kit v.2, Chromium Genome Chip Kit v.2, Chromium i7 Multiplex Kit, and Chromium controller according to manufacturer’s instructions with one modification. Briefly, gDNA was combined with Master Mix, a library of Genome Gel Beads, and partitioning oil to create Gel Bead-in-Emulsions (GEMs) on a Chromium Genome Chip. The GEMs were isothermally amplified. Prior to Illumina library construction, the GEM amplification product was sheared on a Covaris E220 Focused Ultrasonicator to ~350bp then converted to a sequencing library following the 10X standard operating procedure. A total of 679.43 M read pairs were sequenced on an Illumina HiSeqX sequencer, and a *de novo* assembly was constructed with Supernova 2.1.1 (Weisenfeld et al. 2018).

A Chicago library was prepared as described previously (Putnam et al. 2016). Briefly, ~500ng of HMW gDNA was reconstituted into chromatin *in vitro* and fixed with formaldehyde. Fixed chromatin was digested with DpnII, the 5’ overhangs filled in with biotinylated nucleotides, and then free blunt ends were ligated. After ligation, crosslinks were reversed, and the DNA. Purified DNA was treated to remove biotin that was not internal to ligated fragments. The DNA was then sheared to ~350 bp mean fragment size and sequencing libraries were generated using NEBNext Ultra enzymes and Illumina-compatible adapters. Biotin-containing fragments were isolated using streptavidin beads before PCR enrichment of each library. The libraries were sequenced on an Illumina HiSeqX to produce 242 million 2×150 bp paired end reads.

A Dovetail Hi-C library was prepared in a similar manner as described previously (Lieberman-Aiden et al. 2009). For each library, chromatin was fixed in place with formaldehyde in the nucleus and then extracted. Fixed chromatin was digested with DpnII, the 5’ overhangs filled in with biotinylated nucleotides, and then free blunt ends were ligated. After ligation, crosslinks were reversed, and the DNA purified from protein. Purified DNA was treated to remove biotin that was not internal to ligated fragments. The DNA was then sheared to ~350 bp mean fragment size and sequencing libraries were generated using NEBNext Ultra enzymes and Illumina-compatible adapters. Biotin-containing fragments were isolated using streptavidin beads before PCR enrichment of each library. The libraries were sequenced on an Illumina HiSeqX to produce 179 million 2×150 bp paired end reads.

The input *de novo* assembly and Chicago library reads, and Dovetail Hi-C library reads were used as input data for assembly scaffolding with HiRise (Putnam et al. 2016). An iterative analysis was conducted. First, Chicago library sequences were aligned to the draft input assembly using a modified SNAP read mapper (http://snap.cs.berkeley.edu). The separations of Chicago read pairs mapped within draft scaffolds were analyzed by HiRise to produce a likelihood model for genomic distance between read pairs, and the model was used to identify and break putative misjoins, to score prospective joins, and make joins above a threshold. After aligning and scaffolding Chicago data, Dovetail Hi-C library sequences were aligned and scaffolded following the same approach.

### Genome Annotation

The Rio Pearlfish genome was annotated in the NCBI Euakryotic genome annotation pipeline v9.0 (Thibaud-Nissen et al. 2016) and MAKER (Bowman et al. 2017; Campbell et al. 2014; Cantarel et al. 2008) using protein evidence from 15 fish species (Supplementary table 1), and transcriptome evidence from Rio Pearlfish DIII and hatched larvae (Thompson & Ortí 2016). Genome assembly and annotation completeness were analyzed with BUSCOv5 (Simão et al. 2015) and CEGMA (Parra et al. 2007) via the gVolante server (Nishimura et al. 2017, https://gvolante.riken.jp).

### Phylogenetics and Orthology

To confirm species identification, we extracted and concatenated the barcoding marker genes *cox1* and *cytb* from our genome assembly, aligned them with orthologous sequences from all three described *Nematolebias* species (Costa et al. 2014) and inferred a phylogeny partitioned by codon and gene with RAXML (Stamatakis 2014, 2006) with the following parameters: -T 4 -N autoMRE -m GTRCAT -c 25 -p 12345 -f a -x 12345 --asc-corr lewis. We used Orthofinder v2.4.1 (Emms & Kelly 2015) to identify orthologous protein sequences between *N. whitei* and 35 other vertebrates genomes and protein sequences obtained from Cui et al. (2019), Hara et al. (2018), and the longest isoforms of other species available on NCBI RefSeq (Supplementary Table 2) downloaded with orthologr’s retrieve longest isoforms function (Drost et al. 2015).

### Synteny and Genome 3-D Structure

We examined conservation of synteny using genome assemblies and NCBI annotations for Rio Pearlfish, medaka (oryLat2, UCSC), and zebrafish (GCF_000002035.5_GRCz10, NCBI) as input for SynMap in CoGe (Lyons & Freeling 2008). Bwa v 0.7.17 (Li & Durbin 2009) was used to independently map Rio Pearlfish Hi-C read pairs to the genome assembly with the following parameters: bwa mem -A1 -B4 -E50 -L0, and HiC explorer was used to construct a Hi-C matrix with the resulting bam files as follows: hicBuildMatrix --binSize 10000 -- restrictionSequence GATC --danglingSequence GATC. The matrix was corrected via hicCorrectMatrix correct --filterThreshold -1.5 5. The matrix was binned depending on preferred resolution for viewing. Contact maps were visualized with hicPlotMatrix --log1p, and compared to contact maps of syntenic regions in medaka and zebrafish (Nakamura et al. 2021).

### Repeat content and TE landscape

We constructed a species-specific repeat database with Repeat Modeler 2.0.1 (Smit & Hubley 2008). This library as well as vertebrate Repbase annotations (Jurka 2000) (downloaded 15 November 2017), and repeat libraries from platyfish (Schartl et al. 2013), coelacanth (Amemiya et al. 2013), bowfin (Thompson et al. 2021), and gar (Braasch et al. 2016) were combined to annotate repeat elements with Repeat Masker v4.0.5 (Smit et al. 2013). CalcDivergenceFromAlign.pl and createRepeatLandscape.pl in the Repeat Masker package were used to generate a repeat landscape. We compared TE landscape of Rio Pearlfish and those described for other sequenced killifish species (Rhee et al. 2017; Cui et al. 2019; Reichwald et al. 2015; Valenzano et al. 2015).

### Hatching Gene Expression Analysis

We BLASTed the well-studied medaka hatching enzyme genes (*lce* and *hce* paralogs) against the Rio Pearlfish genome to ensure that Pearlfish hatching enzyme genes were not missed by annotation pipelines. We confirmed Pearlfish hatching enzyme gene orthology to those of other teleosts by best BLAST hits to medaka, *Austrofundulus, Kryptolebias,* and *Nothobranchius* metalloprotease genes and inferred gene trees with these sequences (data not shown). Hatching enzyme gene annotations were examined for transcript evidence from DIII embryos (Thompson & Ortí 2016) to identify active *lce* and *hce* gene expression in Rio Pearlfish diapause. We identified a tandem duplication of the low-choriolytic enzyme (*lce.1* and lce.2, Supplementary Text 1) specific to this species and supported by transcript evidence (Thompson & Ortí 2016). We generated an antisense probe of *lce.2* and performed whole mount RNA *in situ* hybridization to identify hatching enzyme expression patterns as marker for the location of HGCs in Rio Pearlfish. Briefly, total RNA was extracted from DIII pearlfish embryos with a Qiagen RNeasy mini plus kit and reverse transcribed with a superscript IV VILO kit from ThermoFisher according to manufacturers’ instructions. The *lce.2* cDNA was amplified from the reverse transcribed template via PCR (see primer sequences in Supplementary Text 1) and inserted into a TOPO TA cloning kit vector (Invitrogen) according to manufacturer’s instructions. Whole mount RNA *in situ* hybridization on manually dechorionated DIII Rio Pearlfish embryos was performed following Deyts et al. (2005) with a 25ug/mL proteinase k digestion treatment for 45min (n=3 embryos), 60min (n=3 embryos), and 90min (n=2 embryos).

## Results and Discussion

### Genome Sequencing and Assembly

We report a high-quality, 1.2 Gb chromosome-level genome assembly of *N.* whitei. The Rio Pearlfish genome assembly consists of 24 chromosomal pseudomolecules represented by 24 superscaffolds and matches the described karyotype (n=24; Von Post, 1965). The assembly has an N50 over 49.98 Mb scaffolds with an L50 of 11 scaffolds (Supplementary Table 1). BUSCO and CEGMA scores for different core gene databases indicate a high-quality assembly with an average of 94% complete BUSCOS and CEGs across all relevant databases (Supplementary Table 1).

### Genome Annotation

The NCBI *Nematolebias whitei* Annotation Release 100.20210725 contains 23,038 genes, with 21,341 protein coding genes, similar to other, chromosomal-level killifish genome assemblies from *Nothobranchius furzeri* and *Kryptolebias marmoratus* (Supplementary Table 1, (Reichwald et al. 2015; Valenzano et al. 2015; Kelley et al. 2016; Rhee et al. 2017). MAKER annotated 26,016 protein coding genes, on par with the NCBI annotation. See Supplementary Table 1 for Rio Pearlfish genome annotation statistics.

### Phylogenetics and Orthology

Our Orthofinder analysis (Fig. 1B, Supplementary Table 2) illustrates the phylogenetic position of Rio Pearlfish among vertebrates and identified 31,317 orthogroups across 36 vertebrate species with 99.2% of Rio Pearlfish genes within orthogroups. We identify 7,287 orthogroups across all species from sharks to human to Rio Pearlfish, highlighting the utility of the Rio Pearlfish genome to connect species with extreme developmental phenotypes to other vertebrates, including traditional vertebrate model species such as mouse, *Xenopus*, zebrafish, etc. We confirm the identity of our genome specimen with barcoding and a molecular phylogeny of *cox1* and *cytb* with its position located within the *N. whitei* clade of *Nematolebias* killifishes (Figure 1B).

### Synteny and Genome 3-D Structure

To confirm the quality of the genome assembly and assess the utility of the chromatin conformation data to interrogate 3-D genome structure and gene regulation, we constructed a Hi-C contact map showing the higher contact frequency within the 24 pearlfish chromosomes (Fig. 1D) than between chromosomes. Using the genome sequence and gene annotations for Rio Pearlfish in synteny comparisons to another atherinomorph teleost, the medaka, (separated by ~85 Million years), and the ostariophysian teleost zebrafish (separated by ~224 million years), we reveal largely conserved synteny for these species across millions of years of evolution (Thompson et al. 2021; Hughes et al. 2018, Fig. 2E,F). We examined a TAD shown to be conserved from zebrafish to medaka (Nakamura et al. 2021) and find high frequency of contacts in the syntenic region between *rasa1a and mctp1a* in Rio Pearlfish liver tissue that strikingly resemble contact maps both in a medaka fibroblast cell line and zebrafish whole embryos (Nakamura et al. 2021, Fig. 1G). Hi-C analyses thus confirm the high-quality of our genome assembly as well as the strikingly conserved nature of 3-D genome interactions across teleost evolution, developmental stages, and among cell and tissue types.

**Figure 2.**
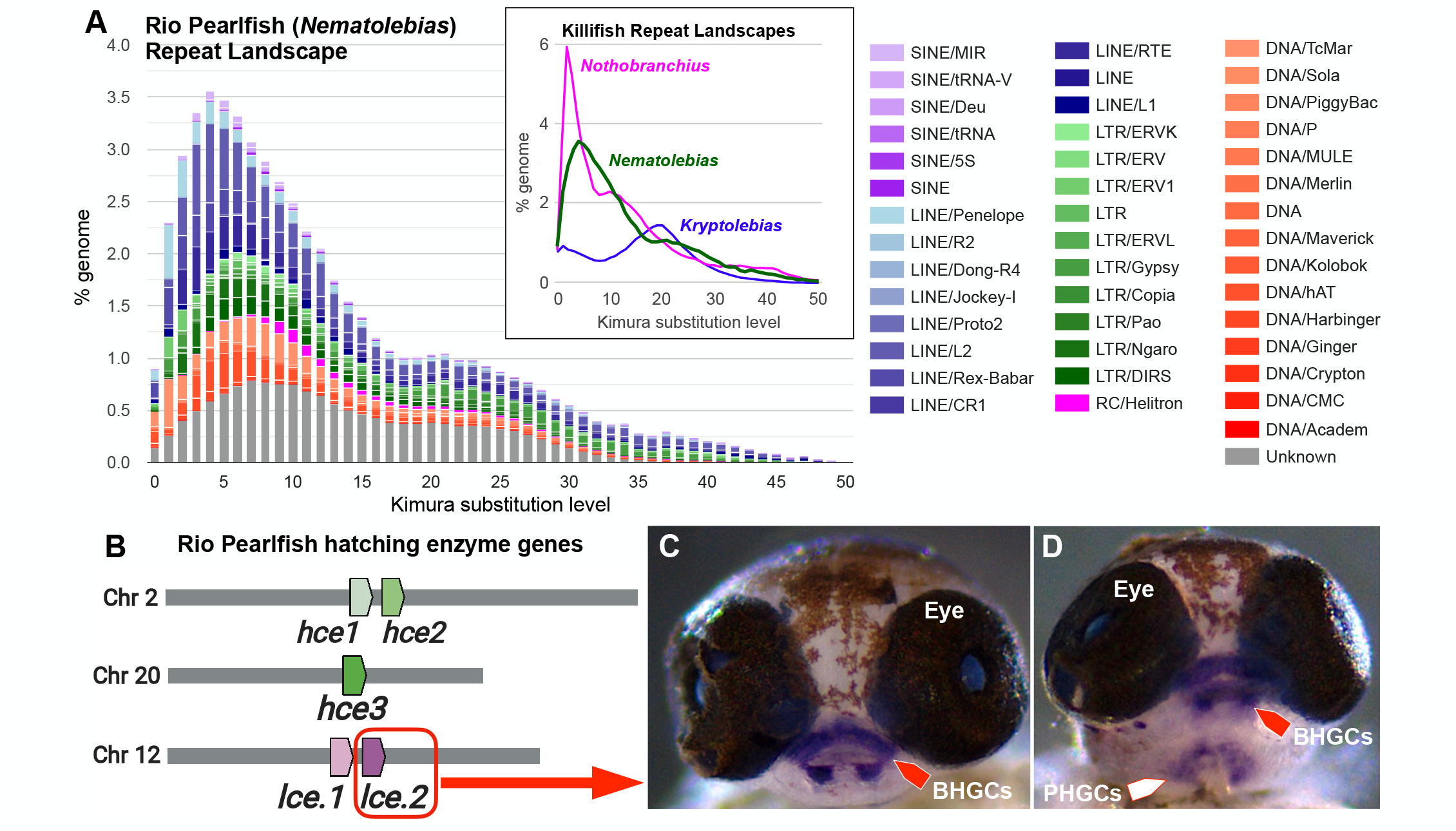
Rio Pearlfish repeat landscape, hatching enzyme genes, and hatching gland location. A.) Repeat landscape of mobile genetic elements in Rio Pearlfish showing a high repeat content with two peaks at Kimura distance 4 and 21. Total transposable element landscape among killifish with independent, recent expansions in the convergent annuals *Nothobranchius* (Cui et al. 2019) and *Nematolebias* (this study) compared to the non-annual *Kryptolebias* (Choi et al. 2020) B.) Locations of five hatching enzyme genes in the Rio Pearlfish genome expressed during DIII. C-D.) Wholemount RNA *in situ* hybridization of *lce.2* in DIII Rio Pearlfish embryos marking hatching gland cells (HGCs) identified in the buccal (BHGCs, red arrows) and pharyngeal (PHGCs, white arrow) cavities.

### Repeat content and transposable element landscape

We find that the Rio Pearlfish genome is highly repetitive with a repeat content of ~57% (Fig. 2A, Supplementary Table 1) which is substantially elevated compared to a non-annual member of the same South American family, *Kryptolebias marmoratus,* with around ~27% (Rhee et al. 2017; Choi et al. 2020). Similarly, African annual *Nothobranchius* killifishes have higher TE repeat content than their non-annual counterparts (Cui et al. 2019). This pattern might be the result of adaptation to extreme environments as animals, fungi, and plants have co-opted TEs for environmental adaptations in harsh conditions (Casacuberta & González 2013; Esnault et al. 2019) and TEs may play roles in vertebrate adaptive radiations (Feiner 2016). Our findings further highlight the expanded repeat content in annual killifish genomes and the Pearlfish genome provides more resources to study the role of mobile DNA in extremophiles.

### Hatching Gene Expression Analysis and hatching gland location

While hatching from the egg is a critical time point during animal development, little is known about its genetic regulation and the integration of environmental cues. Rio Pearlfish is a tractable model for studying hatching regulation since hatching is easily induced in this species by exposing DIII embryos to water (Thompson 2016). Thus, we examined the hatching enzyme gene repertoire and hatching gland location in Pearlfish which reveals five expressed hatching enzyme genes (Fig. 2B, three *hce* and two *lce)* upon mapping DIII mRNA reads from Thompson & Ortí (2016) to the genome. We annotate *hce1* and *hce2* on chromosome 2 (corresponding NCBI GeneIDs: 119423801, 119423789), and *hce3* on chromosome 20 (corresponding NCBI GeneID: 119426643) and the adjacent *lce.1* and *lce.2* genes (corresponding NCBI GeneIDs: 119418488, 119418489) on chromosome 12 that are tandem duplicates (Fig. 2B, Supplementary Text 1). Using whole mount RNA *in situ* hybridization for *lce.2* in DIII embryos, we identify HGC locations in the buccal and pharyngeal cavity in Rio Pearlfish (Fig. 2C,D) similar to HGC localization in medaka (Inohaya, et al. 1995) and the related mummichog or Atlantic killifish (*Fundulus heteroclitus*) (Kawaguchi et al. 2005). These findings will be instrumental in future studies on hatching regulation and hatching physiology among annual and non-annual killifishes.

## Conclusions

Our chromosome-level, dually annotated genome assembly of the Rio Pearlfish provides a valuable comparative genomics resource strengthening the utility of killifishes for studying aging, suspended animation, and response to environmental stress. The Rio Pearlfish is an emerging “in extremo” Eco-Evo-Devo research organism, and this reference genome will be a substrate for future functional genetic and multi-omic approaches exploring how organisms integrate developmental and environmental cues to adapt to extreme environmental conditions in a constantly changing world.

## Acknowledgements

We thank Camilla Peabody for guidance with RNA *in situ* hybridization, Kevin Childs for computational resources, and Françoise Thibaud-Nissen for help integrating the genome into NCBI’s Eukaryotic Annotation Pipeline. This work was supported by the NSF BEACON Center for the Study of Evolution in Action (Cooperative Agreement No. DBI-0939454), project #1233.

## Author Contributions

AWT and IB conceived the project, wrote the manuscript, and acquired funding; genome sequencing and assembly was performed with Dovetail Genomics; MD, HWP, AWT, and IB analyzed hatching enzyme genes; HWP and AWT performed RNA *in situ* hybridization; AWT performed comparative genomic analyses, and genome structure analyses; AWT and IB analyzed repeat content.

## Data Availability

The genome sequence, annotation, and sequence read data are available on NCBI under accession GCA_014905685.2 and Bioproject PRJNA560526. The genome assembly and annotation has also been integrated to the University of California Santa Cruz Genome Browser (https://hgdownload.soe.ucsc.edu/hubs/fish/index.html). The MAKER genome annotation is available on github (https://github.com/AndrewWT?tab=repositories).

---

**Supplementary Table1.**
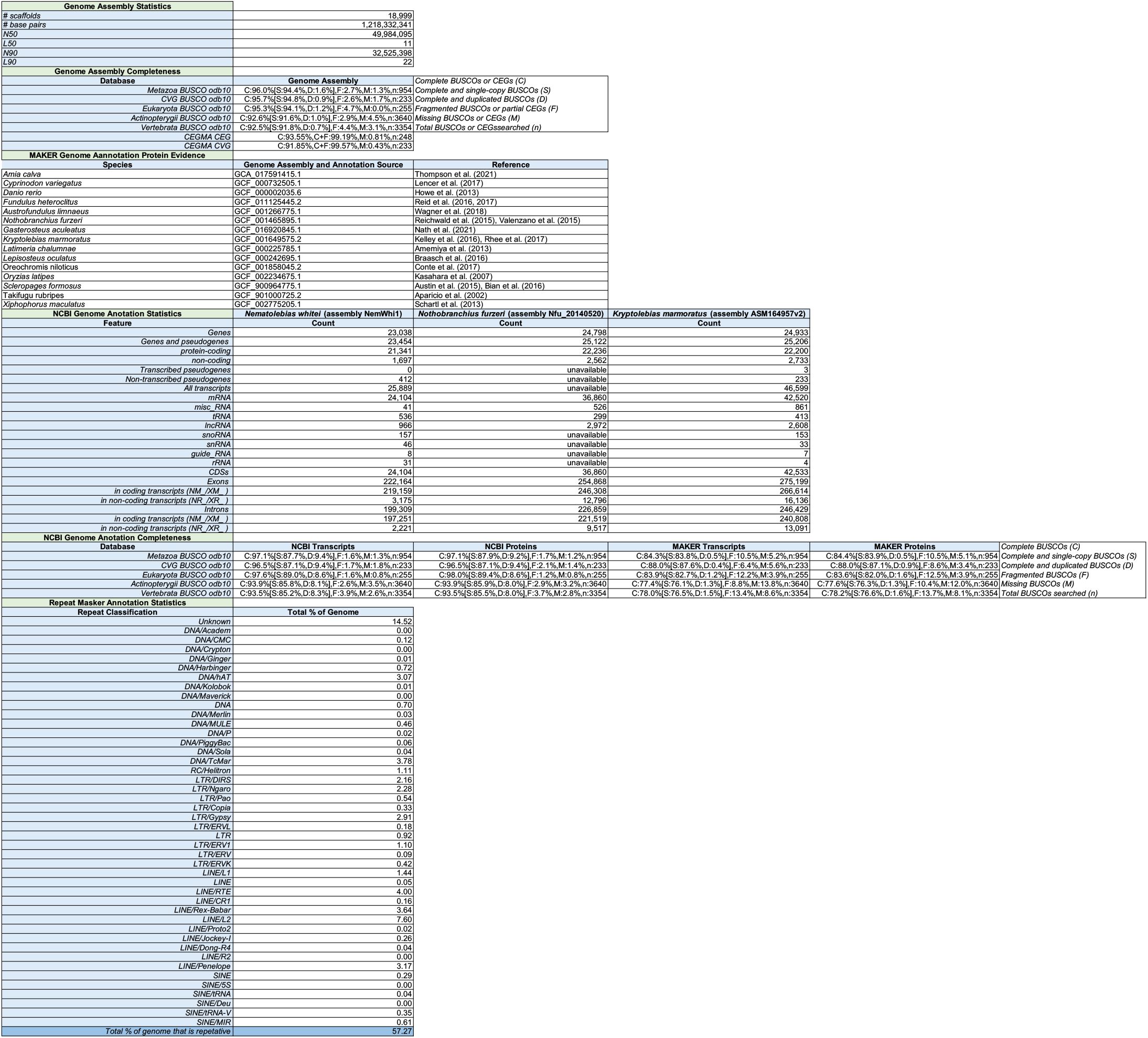
Rio Pearlfish genome assembly and annotation statistics.

**Supplementary Table 2.**
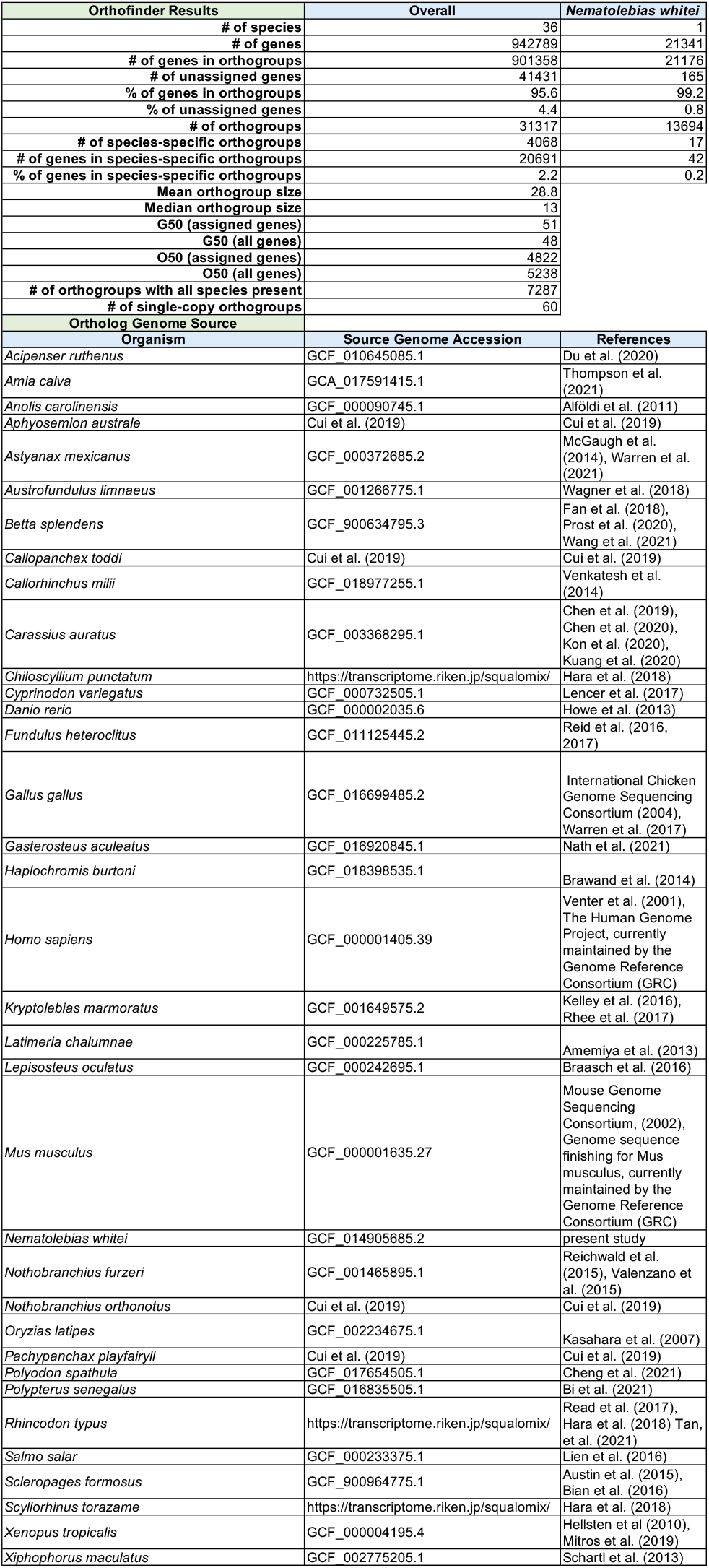
Orthology Statistics.

| Orthofinder Results |  | Overall | *Nematolebias whitei* |
| --- | --- | --- | --- |
| # of species |  | 36 | 1 |
| # of genes |  | 942789 | 21341 |
| # of genes in orthogroups |  | 901358 | 21176 |
| # of unassigned genes |  | 41431 | 165 |
| % of genes in orthogroups |  | 95.6 | 99.2 |
| % of unassigned genes |  | 4.4 | 0.8 |
| # of orthogroups |  | 31317 | 13694 |
| # of species-specific orthogroups |  | 4068 | 17 |
| # of genes in species-specific orthogroups |  | 20691 | 42 |
| % of genes in species-specific orthogroups |  | 2.2 | 0.2 |
| Mean orthogroup size |  | 28.8 |  |
| Median orthogroup size |  | 13 |  |
| G50 (assigned genes) |  | 51 |  |
| G50 (all genes) |  | 48 |  |
| O50 (assigned genes) |  | 4822 |  |
| O50 (all genes) |  | 5238 |  |
| # of orthogroups with all species present |  | 7287 |  |
| # of single-copy orthogroups |  | 60 |  |
| Ortholog Genome Source |  | Source Genome Accession | References |
| *Acipenser ruthenus* |  | GCF_010645085.1 | Du et al. (2020) |
| *Amia calva* |  | GCA_017591415.1 | Thompson et al. (2021) |
| *Anolis carolinensis* |  | GCF_000090745.1 | Alfoldi et al. (2011) |
| *Aphyosemion australe* |  | Cui et al. (2019) | Cui et al. (2019) |
| *Astyanax mexicanus* |  | GCF_000372685.2 | McGaugh et al. (2014), Warren et al. (2021) |
| *Austrofundulus limnaeus* |  | GCF_001266775.1 | Wagner et al. (2018) |
| *Betta splendens* |  | GCF_900634795.3 | Fan et al. (2018), Prost et al. (2020), Wang et al. (2021) |
| *Callopanchax toddi* |  | Cui et al. (2019) | Cui et al. (2019) |
| *Callorhynchus milii* |  | GCF_018977255.1 | Venkatesh et al. (2014) |
| *Carassius auratus* |  | GCF_003368295.1 | Chen et al. (2019), Chen et al. (2020), Kon et al. (2020), Kuang et al. (2020) |
| *Chiloscyllium punctatum* |  | https://transcriptome.riken.jp/squalomix/ | Hara et al. (2018) |
| *Cyprinodon variegatus* |  | GCF_000732505.1 | Lencer et al. (2017) |
| *Danio rerio* |  | GCF_000002035.6 | Howe et al. (2013) |
| *Fundulus heteroclitus* |  | GCF_011125445.2 | Reid et al. (2016, 2017) |
| *Gallus gallus* |  | GCF_016699485.2 | International Chicken Genome Sequencing Consortium (2004), Warren et al. (2017) |
| *Gasterosteus aculeatus* |  | GCF_016920845.1 | Nath et al. (2021) |
| *Haplochromis burtoni* |  | GCF_018398535.1 | Brawand et al. (2014) |
| *Homo sapiens* |  | GCF_000001405.39 | Venter et al. (2001), The Human Genome Project, currently maintained by the Genome Reference Consortium (GRC) |
| *Kryptolebias marmoratus* |  | GCF_001649575.2 | Kelley et al. (2016), Rhee et al. (2017) |
| *Latimeria chalumnae* |  | GCF_000225785.1 | Amemiya et al. (2013) |
| *Lepisosteus oculatus* |  | GCF_000242695.1 | Braasch et al. (2016) |
| *Mus musculus* |  | GCF_000001635.27 | Mouse Genome Sequencing Consortium, (2002), Genome sequence finishing for Mus musculus, currently maintained by the Genome Reference Consortium (GRC) |
| *Nematolebias whitei* |  | GCF_014905685.2 | present study |
| *Nothobranchius furzeri* |  | GCF_001465895.1 | Reichwald et al. (2015), Valenzano et al. (2015) |
| *Nothobranchius orthonotus* |  | Cui et al. (2019) | Cui et al. (2019) |
| *Oryzias latipes* |  | GCF_002234675.1 | Kasahara et al. (2007) |
| *Pachypanchax playfairii* |  | Cui et al. (2019) | Cui et al. (2019) |
| *Polyodon spathula* |  | GCF_017654505.1 | Cheng et al. (2021) |
| *Polypterus senegalus* |  | GCF_016835505.1 | Bi et al. (2021) |
| *Rhincodon typus* |  | https://transcriptome.riken.jp/squalomix/ | Read et al. (2017), Hara et al. (2018) Tan, et al. (2021) |
| *Salmo salar* |  | GCF_000233375.1 | Lien et al. (2016) |
| *Scleropages formosus* |  | GCF_900964775.1 | Austin et al. (2015), Bian et al. (2016) |
| *Scyliorhinus torazame* |  | https://transcriptome.riken.jp/squalomix/ | Hara et al. (2018) |
| *Xenopus tropicalis* |  | GCF_000004195.4 | Hellsten et al. (2010), Mitros et al. (2019) |
| *Xiphophorus maculatus* |  | GCF_002775205.1 | Schartl et al. (2013) |

## Supplementary Text 1

*lce* primers:

Nwh_lce.1_1F: 5’-ATGGACCATAAAGCAAAAGTTTCTCTC-3’

Nwh_lce.1_792R: 5’-CTATTGCTTGTATTTTGAACACTTGT-3’

Nwh_lce.2_1F: 5’-ATGGACCATAAAGCAAAAGTTACTCTT-3’

Nwh_lce.2_825R: 5’-CTATTGCTTGTATTTTGAACAGTTGT-3’

product sizes = 792 and 825 bp, respectively

>lce.1_corresponding_NCBI_GeneID_119418488 ATGGACCATAAAGCAAAAGTTTCTCTCTTGCTCTTGCTGCTTTCAGGCCTCTGCAATGCTCA CACCAGAGATCACTCGAAGAAGGAGGATATTTCTACAACGATCATCAGAATAAACAATGGA ACTTTGGTTCAACAACTGATTGAAGGAGATGTGGTCATTCCAAAAACCAGGAATGCTATGA AGTGCCACACTAAACAATACAGCTGCTTCTGGCCAAAGTCTACCAACGGGAATGTAGAAGT CCCTTTTGTTATAAGTCCCAAGTATGATGACGATGAGAGGAATATAATTCTGACTGCCATGA AAGGCTTTGAACCAAAGACCTGCATTCGCTTTGTTCCTCGTACAAAACAAAGGGCATACCT AAGCATTGAACCAAGATTTGGTTGCTTTTCTTTGCTGGGTCCTACCGGAGAGAAGCAACTC GTGTCTCTGCAGAGAGCTGGCTGCGTGGACAATGGGATCGTCCAACATGAGCTGCTGCAT GCTCTGGGTTTCTACCACGAACACAACCGCAGCGACCGTGACAAGTATGTCAAGATCCACT GGGAAAACATGCATGATGATTTTAAGACCTACTTCAGCAAGATGGATACAGACAATCTCAAT ACCAAATATGACTACTCATCTGTGATGCATTATGGAAAAACTGCCTTTGGAACGAATGGGAA AGAAACCATAACTCCCATCCCCGATCCCAATGTTCCCCTTGGCCAAAGGGTTGGCATGTCT GATATCGACATTGTCAGAGTCAACAGGCTGTACAAGTGTTCAAAATACAAGCAATAG

>lce.2_corresponding_NCBI_GeneID_119418489 ATGGACCATAAAGCAAAAGTTACTCTTTTGCTCTTGCTGCTTTCAGGCCTCTGCAATGCTCA CACTGGAGATCACTCCAAGAAGGAGGATATTTCTACAACGATCATCAGAATGAACAATGGA ACTTTGGTTCAACAACTGATTGAAGGAGATATGCTTATTCCAAAAACCAGGAATGCTTTGAA GTGCCACAATAAACAATACAGCTGCTTCTGGCCAAAGTCTACCAACGGGAATGTAGAAGTC CCTTTTGTTATAAGTCCCAAGTATGATAAAGATGAGAGGAAAACAATTCTGACTGCCATGAA AGGCTTTGAACCAAAGACCTGCATTCGATTTGTTCCTCGTACAAATCAAAGGGCACACCTA AGCCTGGAACCAAAATTTGGTTGCTTTTCTTCTCTGGGACGTGTCGGAGAGAAGCAACTCG TGTCTCTGCAAAGATATGGCTGTGTGGAAAAAGGGATCGTCCAACATGAGCTGCTGCATGC TCTGGGTTTCTACCACGAACACAACCGCAGCGACCGTGACAAGTATGTCAAGATCCACTG GGACAATATGCCAGATATTGTTAAGGTCAACTTCAAAAAAATGGATACAGACAATCTCAACA CCAAATATGACTACTCATCTGTGATGCAATATGGAAAAACTGCCTTTGGAACGGATGGAAA AGAAACCATAACTCCCATCCCCGATCCCAATGTTCCCATCGGCCAAAGAGTGGGCATGTCT GATATTGACATTCTCAGAGTCAACAGGCTGTACAACTGTTCAAAATACAAGCAATAG

## Notes

### Competing Interest Statement

The authors have declared no competing interest.

## References

Alfred Aluru N et al. 2015. Targeted mutagenesis of aryl hydrocarbon receptor 2a and 2b genes in Atlantic killifish (Fundulus heteroclitus). Aquat. Toxicol. 158:192–201. doi: 10.1016/j.aquatox.2014.11.016.

Amemiya C et al. 2013. The African coelacanth genome provides insights into tetrapod evolution. Nature. 496:311–316. doi: 10.1038/nature12027.

Bowman MJ, Pulman JA, Liu TL, Childs KL. 2017. A modified GC-specific MAKER gene annotation method reveals improved and novel gene predictions of high and low GC content in Oryza sativa. BMC Bioinformatics. 18:1–15. doi: 10.1186/s12859-017-1942-z.

Braasch I et al. 2015. A new model army: Emerging fish models to study the genomics of vertebrate Evo-Devo. J. Exp. Zool. Part B Mol. Dev. Evol. 324:316–341. doi: 10.1002/jez.b.22589.

Braasch I et al. 2016. The spotted gar genome illuminates vertebrate evolution and facilitates human-teleost comparisons. Nat. Genet. 48:427–437. doi: 10.1038/ng.3526.

Bravo JI, Nozownik S, Danthi PS, Benayoun BA. 2020. Transposable elements, circular RNAs and mitochondrial transcription in age-related genomic regulation. Development. 147:1–18. doi: 10.1242/dev.175786.

Campbell MS, Holt C, Moore B, Yandell M. 2014. Genome Annotation and Curation Using MAKER and MAKER-P. John Wiley and Sons Inc. doi: 10.1002/0471250953.bi0411s48.

Cantarel BL et al. 2008. MAKER: An easy-to-use annotation pipeline designed for emerging model organism genomes. Genome Res. 18:188–196. doi: 10.1101/gr.6743907.

Carter CA, Wourms JP. 1993. Naturally occurring diblastodermic eggs in the annual fish Cynolebias: Implications for developmental regulation and determination. J. Morphol. 215:301–312. doi: 10.1002/jmor.1052150309.

Casacuberta E, González J. 2013. The impact of transposable elements in environmental adaptation. Mol. Ecol. 22:1503–1517. doi: 10.1111/mec.12170.

Choi BS et al. 2020. The reference genome of the selfing fish Kryptolebias hermaphroditus: Identification of phases I and II detoxification genes. Comp. Biochem. Physiol. - Part D Genomics Proteomics. 35. doi: 10.1016/j.cbd.2020.100684.

Costa WJEM. 2002. The neotropical seasonal fish genus Nematolebias (Cyprinodontiformes Rivulidae Cynolebiatinae) taxonomic revision with description of a new species. Ichthyol. Explor. Freshw. 1:41–52.

Costa WJEM, Amorim PF, Aranha GN. 2014. Species Limits and DNA Barcodes in Nematolebias, a Genus of seasonal Killifishes Threatened with Extinction from the Atlantic Forest of South-Eastern Brazil, with Description of a New Species (Teleostei Rivulidae). Ichthyol. Explor. Freshwaters. 24:225–236.

Cui R et al. 2019. Relaxed Selection Limits Lifespan by Increasing Mutation Load. Cell. doi: 10.1016/j.cell.2019.06.004.

Deyts C, Candal E, Joly JS, Bourrat F. 2005. An automated in situ hybridization screen in the medaka to identify unknown neural genes. Dev. Dyn. 234:698–708. doi: 10.1002/dvdy.20465.

Drost HG, Gabel A, Grosse I, Quint M. 2015. Evidence for active maintenance of phylotranscriptomic hourglass patterns in animal and plant embryogenesis. Mol. Biol. Evol. 32:1221–1231. doi: 10.1093/molbev/msv012.

Emms DM, Kelly S. 2015. OrthoFinder: solving fundamental biases in whole genome comparisons dramatically improves orthogroup inference accuracy. Genome Biol. 16:1–14. doi: 10.1186/s13059-015-0721-2.

Esnault C, Lee M, Ham C, Levin HL. 2019. Transposable element insertions in fission yeast drive adaptation to environmental stress. Genome Res. 29:85–95. doi: 10.1101/gr.239699.118.

Feiner N. 2016. Accumulation of transposable elements in hox gene clusters during adaptive radiation of anolis lizards. Proc. R. Soc. B Biol. Sci. 283. doi: 10.1098/rspb.2016.1555.

Furness AI, Reznick DN, Springer MS, Meredith RW. 2015. Convergent evolution of alternative developmental trajectories associated with diapause in African and South American killifish. Proc. R. Soc. B Biol. Sci. 282:20142189.

Hara Y et al. 2018. Shark genomes provide insights into elasmobranch evolution and the origin of vertebrates. Nat. Ecol. Evol. 2:1761–1771. doi: 10.1038/s41559-018-0673-5.

Harel I et al. 2015. A platform for rapid exploration of aging and diseases in a naturally short-lived vertebrate. Cell. 160:1013–1026. doi: 10.1016/j.cell.2015.01.038.

Hong CS, Saint-Jeannet JP. 2014. Xhe2 is a member of the astacin family of metalloproteases that promotes Xenopus hatching. Genesis. 52:946–951. doi: 10.1002/dvg.22841.

Hughes LC et al. 2018. Comprehensive phylogeny of ray-finned fishes (Actinopterygii) based on transcriptomic and genomic data. Procedings Natl. Acad. Sci.

Inohaya K et al. 1997. Species-dependent migration of fish hatcing gland cells that commonly express astacin-like proteases in common. Dev. Growth, Differ. 39:191–197.

Inohaya K, Yasumasu S, Ishimaru M, Ohyama A, Luchi I, et al. 1995. Temporal and spatial patterns of gene expression for the hatching enzyme in the teleost embryo, Oryzias latipes. Dev. Biol. 171:374–385.

Inohaya K, Yasumasu S, Ishimaru M, Ohyama A, Iuchi I, et al. 1995. Temprora and spatial patterns of gene expression for the hatching enzyme in the teleost embryo, Oryzias latipes.pdf. Dev. Biol. 171:374–385.

Jurka J. 2000. Repbase update: a database and an electronic journal of repetitive elements. Trends Genet. 16:418–420. http://www.girinst.org.

Kawaguchi M et al. 2010. Intron-loss evolution of hatching enzyme genes in Teleostei. BMC Evol. Biol. 10:1–10. doi: 10.1186/1471-2148-10-260.

Kawaguchi M et al. 2005. Purification and gene cloning of Fundulus heteroclitus hatching enzyme: A hatching enzyme system composed of high choriolytic enzyme and low choriolytic enzyme is conserved between two different teleosts, Fundulus heteroclitus and medaka Oryzias latipes. FEBS J. 272:4315–4326. doi: 10.1111/j.1742-4658.2005.04845.x.

Kawaguchi M, Yasumasu S, Hiroi J. 2006. Evolution of teleostean hatching enzyme genes and their paralogous genes. Dev. Genes Evol. 216:769–784. doi: 10.1007/s00427-006-0104-5.

Kelley JL et al. 2016. The genome of the self-fertilizing mangrove rivulus fish, *Kryptolebias marmoratus*?: a model for studying phenotypic plasticity and adaptations to extreme environments. Genome Biol. Evol. evw145. doi: 10.1093/gbe/evw145.

Korwin-Kossakowski M. 2012. Fish hatching strategies: A review. Rev. Fish Biol. Fish. 22:225–240. doi: 10.1007/s11160-011-9233-7.

Krefting J, Andrade-Navarro MA, Ibn-Salem J. 2018. Evolutionary stability of topologically associating domains is associated with conserved gene regulation. BMC Biol. 16:1–12. doi: 10.1186/s12915-018-0556-x.

Li H, Durbin R. 2009. Fast and Accurate Short Read Alignment with Burrows-Wheeler Transform. Bioinformatics. 25:1754–1760. doi: 10.1093/bioinformatics/btp324.

Lieberman-Aiden E et al. 2009. Comprehensive Mapping of Long-Range Interactions Reveals Folding Principles of the Human Genome. Science (80-.). 326:285–289. doi: 10.1126/science.1178746.

Lyons E, Freeling M. 2008. How to usefully compare homologous plant genes and chromosomes as DNA sequences. Plant J. 53:661–673. doi: 10.1111/j.1365-313X.2007.03326.x.

Myers GS. 1952. Annual Fishes. Aquarium J. 23:125–141.

Myers GS. 1942. Studies on South American freshwater fishes I. Stanford Ichthyol. Bull. 2:89–114. doi: 10.1108/eb025354.

Nagasawa T et al. 2016. Evolutionary Changes in the Developmental Origin of Hatching Gland Cells in Basal Ray-Finned Fishes. Zoolog. Sci. 33:272–281. doi: 10.2108/zs150183.

Nakamura R et al. 2021. CTCF looping is established during gastrulation in medaka embryos. Genome Res. 31:968–980. doi: 10.1101/gr.269951.120.

Nishimura O, Hara Y, Kuraku S. 2017. GVolante for standardizing completeness assessment of genome and transcriptome assemblies. Bioinformatics. 33:3635–3637. doi: 10.1093/bioinformatics/btx445.

Parra G, Bradnam K, Korf I. 2007. CEGMA: A pipeline to accurately annotate core genes in eukaryotic genomes. Bioinformatics. 23:1061–1067. doi: 10.1093/bioinformatics/btm071.

Von Post A. 1965. Vergleichende Untersuchungen der Chromosomenzahlen bei Süßwasser Teleosteern. Zeitschrift für Zool. Syst. und Evol. 47–93.

Putnam NH et al. 2016. Chromosome-scale shotgun assembly using an in vitro method for long-range linkage. Genome Res. 26:342–350. doi: 10.1101/gr.193474.115.Freely.

Reichwald K et al. 2015. Insights into Sex Chromosome Evolution and Aging from the Genome of a Short-Lived Fish. Cell. 163:1527–1538.

Rhee JS et al. 2017. Diversity, distribution, and significance of transposable elements in the genome of the only selfing hermaphroditic vertebrate Kryptolebias marmoratus. Sci. Rep. 7. doi: 10.1038/srep40121.

Ruijter JM. 1987. Development and aging of the teleost pituitary: qualitative and quantitative observations in the annual cyprinodont Cynolebias whitei. Anat. Embryol. (Berl). 175:379–386. doi: 10.1007/BF00309851.

Ruijter JM, Creuwels LAJM. 1988. The ultrastructure of prolactin cells in the annual cyprinodont Cynolebias whitei during its life cycle. Cell Tissue Res. 253:477–483.

Ruijter JM, Kemenade JAMVan, Bonga SEW. 1984. Environmental influences on prolactin cell development in the cyprinodont fish, Cynolebias whitei. Cell Tissue Res. 238:595–600.

Sanger TJ, Rajakumar R. 2019. How a growing organismal perspective is adding new depth to integrative studies of morphological evolution. Biol. Rev. 94:184–198. doi: 10.1111/brv.12442.

Sano K, Kawaguchi M, Watanabe S, Yasumasu S. 2014. Neofunctionalization of a duplicate hatching enzyme gene during the evolution of teleost fishes. BMC Evol. Biol. 14:221. doi: 10.1186/s12862-014-0221-0.

Schartl M et al. 2013. The genome of the platyfish, Xiphophorus maculatus, provides insights into evolutionary adaptation and several complex traits. Nat Genet. 45:567–572. doi: 10.1038/ng.2604.

Schoots AFM, Ruijter JM, van Kemenade JAM, Denucé JM. 1983. Immunoreactive prolactin in the pituitary gland of cyprinodont fish at the time of hatching. Cell Tissue Res. 233:611–618. doi: 10.1007/BF00212228.

Shimizu D et al. 2014. Comparison of Hatching Mode in Pelagic and Demersal Eggs of Two Closely Related Species in the Order Pleuronectiformes. Zoolog. Sci. doi: 10.2108/zs140018.

Simão FA, Waterhouse RM, Ioannidis P, Kriventseva E V., Zdobnov EM. 2015. BUSCO: Assessing genome assembly and annotation completeness with single-copy orthologs. Bioinformatics. 31:3210–3212. doi: 10.1093/bioinformatics/btv351.

Simpson BRC. 1979. The phenology of annual killifish. Symp Zool Soc Lon. 44:243–261.

Smit AFA, Hubley R. 2008. RepeatModeler Open-1.0. http://www.repeatmasker.org.

Smit AFA, Hubley R, Green P. 2013. RepeatMasker Open-4.0. <http://www.repeatmasker.org>.

Stamatakis A. 2006. RAxML-VI-HPC: Maximum likelihood-based phylogenetic analyses with thousands of taxa and mixed models. Bioinformatics. 22:2688–2690. doi: 10.1093/bioinformatics/btl446.

Stamatakis A. 2014. RAxML version 8: A tool for phylogenetic analysis and post-analysis of large phylogenies. Bioinformatics. 30:1312–1313. doi: 10.1093/bioinformatics/btu033.

Thibaud-Nissen F et al. 2016. P8008 The NCBI Eukaryotic Genome Annotation Pipeline. J. Anim. Sci. 94:184.

Thompson AW et al. Deterministic shifts in molecular evolution correlate with convergence to annualism in killifishes.

Thompson AW et al. 2021. The bowfin genome illuminates the developmental evolution of ray-finned fishes. Nat. Genet.

Thompson AW, Furness AI, Stone C, Rade C, Ortí G. 2017. Microanatomical diversification of the zona pellucida in aplochelioid killifishes. J. Fish Biol. 91:126–143.

Thompson AW, Ortí G. 2016. Annual killifish transcriptomics and candidate genes for metazoan diapause. Mol. Biol. Evol. 33:2391–2395. doi: 10.1093/molbev/msw110.

Valenzano DR et al. 2015. The African Turquoise Killifish Genome Provides Insights into Evolution and Genetic Architecture of Lifespan. Cell. 163:1539–1554. doi: 10.1016/j.cell.2015.11.008.

Valenzano DR, Sharp SC, Brunet A. 2011. Transposon-Mediated Transgenesis in the Short-Lived African Killifish Nothobranchius furzeri, a Vertebrate Model for Aging. G3 (Bethesda). 1:531–8. doi: 10.1534/g3.111.001271.

Wagner JT et al. 2018. The genome of Austrofundulus limnaeus offers insights into extreme vertebrate stress tolerance and embryonic development. BMC Genomics. 19:1–21.

Weisenfeld NI, Kumar V, Shah P, Church DM, Jaffe DB. 2018. Corrigendum: Direct determination of diploid genome sequences. Genome Res. 28:757–767. doi: 10.1101/gr.235812.118.

Wourms JP. 1967. Annual Fishes. In: Methods in Developmental Biology. Thomas and Crowell Company: New York pp. 123–137.

Wourms JP. 1972a. The Developmental Biology of Annual Fishes I. Stages in the Normal Development of Austrofundulus Myersi Dahl. J. Exp. Zool. 182:143–168.

Wourms JP. 1972b. The Developmental Biology of Annual Fishes II. Naturally Occurring Dispersion and Reaggregation of Blastomeres During the Development of Annual Fish Eggs. J. Exp. Zool. 182:169–200.

Wourms JP. 1972c. The Developmental Biology of Annual Fishes III. Pre-embryonic and Embryonic Diapause of Variable Duration in the Eggs of Annual Fishes. J. Exp. Zool. 182:389–414.

Yamagami K. 1988. Mechanisms of Hatching in Fish. In: Fish Physiology.Vol. XIA pp. 447–499. doi: 10.1163/157338207X231404.

Yasumasu S et al. 1992. Isolation of cDNAs for LCE and HCE, two constituent proteases of the hatching enzyme of Oryzias latipes, and concurrent expression of their mRNAs during development. Dev. Biol. 153:250–258. doi: 10.1016/0012-1606(92)90110-3.

